# Molecular basis for the distinct cellular functions of the Lsm1-7 and Lsm2-8 complexes

**DOI:** 10.1101/2020.04.22.055376

**Authors:** Eric J. Montemayor, Johanna M. Virta, Samuel M. Hayes, Yuichiro Nomura, David A. Brow, Samuel E. Butcher

## Abstract

Eukaryotes possess eight highly conserved Lsm (like Sm) proteins that assemble into circular, heteroheptameric complexes, bind RNA, and direct a diverse range of biological processes. Among the many essential functions of Lsm proteins, the cytoplasmic Lsm1-7 complex initiates mRNA decay, while the nuclear Lsm2-8 complex acts as a chaperone for U6 spliceosomal RNA. It has been unclear how these complexes perform their distinct functions while differing by only one out of seven subunits. Here, we elucidate the molecular basis for Lsm-RNA recognition and present four high-resolution structures of Lsm complexes bound to RNAs. The structures of Lsm2-8 bound to RNA identify the unique 2′,3′ cyclic phosphate end of U6 as a prime determinant of specificity. In contrast, the Lsm1-7 complex strongly discriminates against cyclic phosphates and tightly binds to oligouridylate tracts with terminal purines. Lsm5 uniquely recognizes purine bases, explaining its divergent sequence relative to other Lsm subunits. Lsm1-7 loads onto RNA from the 3′ end and removal of the Lsm1 C-terminal region allows Lsm1-7 to scan along RNA, suggesting a gated mechanism for accessing internal binding sites. These data reveal the molecular basis for RNA binding by Lsm proteins, a fundamental step in the formation of molecular assemblies that are central to eukaryotic mRNA metabolism.

## Introduction

The Lsm/Sm proteins are an ancient family of RNA binding proteins found in all three domains of life and have a wide range of biological functions. They are named after the autoimmune patient serum that led to their discovery (1). The Sm family includes the Sm, Lsm (“Like Sm”), and bacterial Hfq proteins (2). These proteins share a conserved “Sm fold” consisting of an N-terminal alpha helix followed by five anti-parallel beta strands that form small beta barrels that assemble into ring-shaped hexamers or heptamers. The eukaryotic Sm proteins form heteroheptamers that interact with the major spliceosomal snRNAs U1, U2, U4 and U5, the minor spliceosomal snRNAs U4^atac^, U11, U12, and telomerase RNA. The Sm-like archaeal proteins (SmAPs) are homologous to the eukaryotic Sm proteins, but their biological roles are less well understood (3). The eukaryotic Lsm proteins form at least four different 6- or 7-subunit complexes (2). In addition, a hybrid complex containing Lsm10, Lsm11, and five Sm proteins is essential for 3’ end processing of histone mRNAs (4,5).

The Lsm1-7 complex is cytoplasmic and mediates messenger-RNA (mRNA) decay, a major post-transcriptional mechanism for regulating gene expression (6-9). Binding of the Lsm1-7 complex to mRNA is a key event in the decay pathway as it also binds the protein Pat1 (10-12). Pat1 then recruits a complex consisting of the decapping enzyme Dcp2 and its activators Dcp1, Edc1 and Edc2 (13). After decapping, the 5′-3′ exoribonuclease Xrn1 degrades the mRNA. Structures have been determined for the isolated Lsm1-7 complex (11,14), and Lsm1-7 with a C-terminally truncated Lsm1 bound to the C-terminal domain of Pat1 (11,15). However, no structures are available for Lsm1-7 bound to RNA, despite the central importance of this interaction in the mRNA decay pathway. In addition to activating mRNA decapping, Lsm1-7 has many other functions including formation of phase-separated processing bodies (P-bodies) (16,17), protecting 3′ ends of mRNA from 3′-5′ degradation by the exosome (18,19), stabilizing specific RNAs during starvation and autophagy (20), suppressing translation of stress-activated RNAs during osmotic shock (21), and promoting translation of viral RNAs (22). Human and *S. pombe* Lsm1-7 complexes bind tightly to oligouridylate (hereafter, oligoU) RNAs (12,23). In contrast, the Lsm1-7-Pat1 complex binds tightly to oligoadenylate (hereafter, oligoA) RNAs (10,12,24). The molecular basis for these interactions is unknown.

The Lsm2-8 complex shares 6 out of 7 subunits with Lsm1-7, localizes in the nucleus, and binds the 3’ ends of U6 and U6^atac^ snRNAs (25-28). U6 snRNA is transcribed by RNA polymerase III, which terminates transcription after synthesis of an oligoU tail at the end of U6 snRNA (29). This tail can then be elongated by the enzyme Tutase (29). Finally, U6 snRNA is processed by the 3′-5′ exoribonuclease Usb1 (29), resulting in a 2′,3′ cyclic phosphate in most organisms (30) (Figure 1). In addition, Lsm2-8 mediates nuclear mRNA decay (31). In the case of *S. pombe*, Lsm2-8 is also known to play an important role in telomerase biogenesis (32).

**Figure 1.**
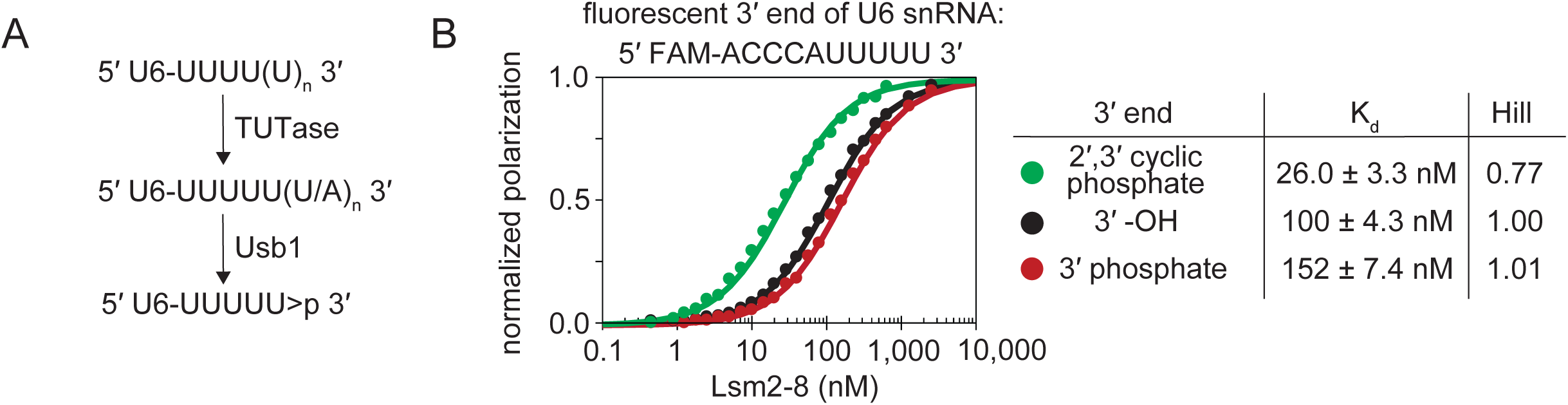
3’-processing of U6 snRNA alters its recognition by *S. pombe* Lsm2-8. A) Processing of U6 by Usb1 leaves a 3′ cyclic phosphate group on U6, denoted as “>p”. B) *In vitro* binding assays show that *S. pombe* Lsm2-8 preferentially binds RNA that has been processed by Usb1. All binding curves and K_d_ determinations in this work were performed with a restrained Hill coefficient of 1. For reference, the non-restrained Hill coefficients, which are close to 1, are shown in the figure.

The molecular basis for the Lsm2-8 interaction with U6 snRNA terminating in a cyclic phosphate has yet to be elucidated, despite existing cryo-EM structures of the human U4/U6.U5 tri-snRNP (33) and pre-B complex (34), both of which contain the Lsm2-8 complex bound to U6 snRNA. In these structures, the U6 snRNA-Lsm2-8 complex can only be resolved to resolutions of ∼10 Å, due to its location on the surface of these complexes and/or dynamic motions. The *S. cerevisiae* Lsm2-8 structure has been determined (25,26) but is unlikely to be representative of most eukaryotes due to significant differences in Lsm8 protein sequence and U6 snRNA post-transcriptional modification (35). Budding yeasts have Usb1 enzymes that open the 2′,3′ cyclic phosphate to produce a 3′ phosphate, and their Lsm8 proteins have evolved a unique C-terminal extension in order to interact with this 3′ phosphate through electrostatic interactions (25,35). In contrast, the Lsm8 proteins of most eukaryotes, from *S. pombe* to humans, do not have this Lsm8 extension and have U6 snRNAs with terminal 2′,3′ cyclic phosphates. The relative specificity of the Lsm2-8 interaction with U6 versus other RNAs is also not well understood.

Here, we investigate the binding properties of Lsm1-7 and Lsm2-8 complexes from *S. pombe*. We show that Lsm2-8 specifically recognizes the terminal 2′,3′ cyclic phosphate while still retaining the ability to bind unmodified RNA. We reveal the molecular basis for Lsm2-8 binding by determining high-resolution structures of Lsm2-8 complexes bound to two RNAs, one that is unmodified and one that has a 2′,3′ cyclic phosphate. Next, we describe the RNA binding properties of the Lsm1-7 complex utilizing a panel of RNAs of varying lengths and sequences. We report two high-resolution structures of Lsm1-7 bound to high affinity RNA targets that explain the molecular basis of Lsm1-7-RNA recognition. We demonstrate that Lsm1-7 loads unidirectionally onto RNA from the 3′ end, using a binding mechanism in which specificity is bestowed by the Lsm1 C-terminal region. Removal of this region enables 3′-5′ scanning and leads to a model for allosteric regulation of mRNA decay. In summary, the RNA binding activities of Lsm1-7 and Lsm2-8 are remarkably distinct despite similar quaternary structures that differ by only one subunit.

## Materials and Methods

### Production of recombinant Lsm2-8

Codon optimized open reading frames (Genscript) for *S. pombe* Lsm proteins were cloned into modified variants of the pQLink expression system (36). In the initial single open reading frame plasmids, Lsm2, Lsm5 and Lsm8 lacked purification tags. Lsm3 and Lsm6 harbored N-terminal glutathione S-transferase and maltose-binding tags, respectively, with an intervening tobacco etch virus (TEV) protease site. Lsm4 and Lsm7 harbored non-cleavable strep-II and hexahistidine tags, respectively, at their C-termini. After cloning individual ORFs into individual plasmids, the multi-ORF expression system was assembled into a single plasmid through ligation independent cloning (36) to yield a final plasmid with ORFs assembled in order Lsm6, Lsm3, Lsm2, Lsm8, Lsm4, Lsm7 and Lsm5. The sequence of the final plasmid was confirmed by Sanger sequencing.

The resulting *S. pombe* Lsm2-8 expression plasmid was transformed into *E. coli* BL21(DE3) STAR pLysS cells (Invitrogen # C602003). Cells were grown in terrific broth (RPI # T15100-5000.0) containing ampicillin with shaking at 37 °C until the OD_600_ was approximately 2, at which point isopropyl β-D-1-thiogalactopyranoside was added to a final concentration of 1 mM and the cells were allowed to continue growing with shaking at 37 °C for another 24 hours, at which point the OD_600_ was approximately 16. Cells were harvested by centrifugation and the resulting cell pellet was resuspended in 0.025 volumes of buffer A (500 mM NaCl, 50 mM HEPES acid, 50 mM imidazole base, 10 % glycerol, 1 mM TCEP, final pH ∼ 7) per original volume of cell culture (*i.e*., 100 mL of resuspension buffer per 4 L of original cell culture). Protease inhibitors (EMD Millipore # 539137-10VL) were added, as were DNaseI (Sigma # DN25-100MG) and lysozyme (Sigma # 62971-10G-F) at a final concentration of 0.02 and 0.5 mg/mL, respectively. The resuspended cells were subjected to a single freeze-thaw cycle prior to sonication and clearance of cell debris by centrifugation.

The soluble fraction was first purified by immobilized metal affinity chromatography (Qiagen # 30230) using gravity flow at room temperature. Buffer A was used for washing and buffer A supplemented with 500 mM imidazole pH ∼ 7 was used for step elution. The pooled eluate was then dialyzed overnight at 4 °C with 20 kDa MWCO membranes (Pierce # 66012) into buffer B (100 mM NaCl, 10 mM HEPES acid, 10 mM sodium HEPES, 10 % glycerol, 1 mM TCEP-HCl, pH ∼ 7.0). Centrifugation was used to remove any precipitation that formed during dialysis. Soluble protein was further purified via glutathione agarose chromatography (GenScript # L00206) with gravity flow at room temperature in fresh buffer B. Step elution was performed in buffer B supplemented with 50 mM HEPES acid, 50 mM sodium HEPES and 10 mM reduced glutathione. One milligram of TEV protease was added to the eluate, which was then dialyzed overnight as above, but at room temperature against 1 L of buffer C (150 mM NaCl, 50 mM tris base, 50 mM tris-HCl, 1 mM trisodium EDTA, 1 mM TCEP-HCl, 1 mM sodium azide, pH ∼ 8.0). Dialyzed protein was then purified using gravity flow chromatography at room temperature on Strep-Tactin agarose (IBA GmBH # 2-1208-025) in fresh buffer C, with step elution in buffer C supplemented with 2.5 mM desthiobiotin. The resulting eluate was manually applied at room temperature to a 1 mL HiTrapQ anion exchange column (GE Healthcare # 29051325) that had been pre-equilibrated in buffer B. The column was then attached to an AKTA chromatography system at 4 °C and elution accomplished by applying a linear gradient of NaCl up to 2 M in buffer B. The Lsm2-8 complex desorbed from the column at approximately 250 mM NaCl. Peak fractions were collected, pooled, and the concentration of the purified complex was determined by UV absorbance and the estimated molar extinction coefficient of 46,300 M^-1^cm^-1^ at 280 nm (37). Final purity of the Lsm2-8 complex was determined by 20 % (29:1) tris-tricine SDS-PAGE. The presence of all protein components was further confirmed by electrospray ionization mass spectrometry at the UW-Madison Biotechnology Center Mass Spectrometry Facility, where the N-terminal methionine of Lsm5 and Lsm8 was found to be missing, and Lsm7 lacked either 1 or 2 N-terminal residues. Protein samples were either stored at 4 °C or frozen as 100 uL aliquots in liquid nitrogen before long-term storage at −80 °C.

### Production of recombinant Lsm1-7

The Lsm1-7 complex was produced as described for Lsm2-8, with the following exceptions. The multi-ORF expression system was assembled into a single plasmid through ligation independent cloning as described for Lsm2-8, with the ORFs assembled in order Lsm6, Lsm3, Lsm2, Lsm1, Lsm4, Lsm7 and Lsm5. Lsm7 lacked a purification tag.

Protein expression, purification and storage was performed as with the above Lsm2-8 complex, with exception that Strep-Tactin chromatography was not used and instead the eluate from glutathione agarose chromatography was loaded directly onto a HiTrapQ column after overnight incubation with TEV protease at room temperature. Peak fractions were collected, pooled, and the concentration of the purified complex was determined by UV absorbance and the estimated molar extinction coefficient at 280 nm (37). Electrospray ionization mass spectrometry showed that all protein chains were full-length with the exception of Lsm5 and Lsm7, which lacked 1 and 1-2 residues from their N-termini, respectively.

In order to prepare the Lsm5-N66A/N68A or Lsm1 C-terminal truncation mutants of Lsm1-7, a modified inverse PCR mutagenesis protocol was employed on the multi-ORF Lsm1-7 expression plasmid. PCR amplification was performed in “GC” optimized Pfusion buffer (NEB # M0532S) using a gradient of annealing temperatures and oligonucleotides that anneal only within the unique open reading frame regions of each Lsm locus. Amplicons of correct length were gel purified prior to phosphorylation and ligation. All mutant clones were confirmed to be of correct length and sequence by analytical restriction enzyme digestion and Sanger sequencing.

### Production of recombinant Prp24

A codon optimized open reading frame (Genscript) for *S. pombe* Prp24 was cloned into a modified variant of plasmid pET3a, encoding an octahistidine tag, biotin acceptor peptide sequence (38), and tobacco etch virus (TEV) protease site located upstream of the Prp24 ORF. The sequence of the final plasmid was confirmed by Sanger sequencing.

The *S. pombe* Prp24 was produced as described above for the Lsm proteins with the following exceptions. Cells were harvested by centrifugation and the resulting cell pellet was resuspended in 0.1 volumes of buffer A (500 mM NaCl, 50 mM HEPES acid, 50 mM imidazole base, 10 % glycerol, 1 mM TCEP, final pH ∼ 7) per original volume of cell culture (ie. 100 mL of resuspension buffer per 1 L or original cell culture).

Soluble protein was manually applied at room temperature to a 5 mL HiTrapQ anion exchange column (GE Healthcare # 29051325) that had been pre-equilibrated in buffer B. The column was then attached to an AKTA chromatography system at 4 °C and elution accomplished by applying a linear gradient of NaCl up to 2 M in buffer B. The peak fractions were collected, pooled, diluted two-fold against fresh buffer B and then manually applied at room temperature to a 5 mL HiTrap Heparin cation exchange column (GE Healthcare # 17040701) that had been pre-equilibrated in buffer B. The column was then attached to an AKTA chromatography system at 4 °C and elution accomplished by applying a linear gradient of NaCl up to 2 M in buffer B. The peak fractions were pooled and the final concentration of protein was determined by UV absorbance and the estimated molar extinction coefficient of 114,600 M^-1^cm^-1^ at 280 nm (37). Protein samples at approximately 6 mg/mL were either stored at 4 °C, or frozen as 100 uL aliquots in liquid nitrogen before long-term storage at −80 °C.

### Synthesis and purification of RNA

*In vitro* transcription with T7 RNA polymerase was used to synthesize mature *S. pombe* U6 nucleotides 1-100 (39) from a linearized pUC57 plasmid template harboring a T7 promoter (TTCTAATACGACTCACTATA) and a minimal 56 nucleotide HDV ribozyme “drz-Mtgn-3” (40) to ensure a homogenous 3′ end with a cyclic phosphate. A 20 mL transcription reaction contained ∼ 0.25 mg/mL linearized plasmid template, ∼ 0.5 mg/mL T7 RNA polymerase, 5 mM each of ATP, GTP, CTP and UTP; 100 mM tris, 100 mM tris-HCl, 40 mM MgCl_2_, 5 mM DTT, 1 mM spermidine trihydrochloride and 0.01 % (v/v) Triton X-100. Synthesis of RNA was performed at 37 °C for 2 hours. Trisodium EDTA pH 8.0 was then added to a final concentration of 100 mM to halt transcription and solubilize accumulated magnesium pyrophosphate. The transcription was then concentrated to ∼ 400 uL with 10 kDa MWCO spin filters (Amicon # UFC901008) prior to addition of 6 mL of 100 % formamide and subsequent electrophoresis on a denaturing 10 % (19:1) polyacrylamide gel containing 8 M urea, 100 mM tris base, 100 mM boric acid and 1 mM EDTA acid. Full-length U6 RNA was identified by UV shadowing, extracted by scalpel, and removed from the gel matrix by passive diffusion overnight at room temperature with gently shaking into a solution containing 300 mM sodium acetate, 50 mM HCl, 1 mM EDTA, 1 mM sodium azide, pH ∼ 5.6. Soluble RNA was separated from solid acrylamide by filtration through 0.22 micron filters and then manually applied to a 5 mL HiTrapQ column (GE Healthcare # 29051325) that had been equilibrated in buffer D (300 mM NaCl, 10 mM KH_2_PO_4_, 10 mM K_2_HPO_4_, 1 mM EDTA acid, 1 mM sodium azide, pH ∼ 7). Bound RNA was washed with 20 mL of buffer D and then eluted in a single step with buffer D adjusted to 2 M NaCl. Eluted RNA was pooled and concentration and buffer exchange were accomplished by three iterations of ten-fold concentration (Amicon # UFC501008) and re-dilution against buffer E (100 mM KCl, 20 mM bis-tris, 10 mM HCl, 1 mM EDTA, 1 mM sodium azide, pH ∼ 6.5). The final RNA concentration was estimated using UV absorption and an anticipated extinction coefficient of 1,223,990 M^-1^cm^-1^ at 260 nm (41).

In order to generate a pentauridylate RNA with a 3′ cyclic phosphate for co-crystallization with *S. pombe* Lsm2-8, an RNA oligo with sequence 5′-UUUUUA-3′ was purchased from Integrated DNA Technologies (IDT) and treated with human Usb1, which rapidly removes 3′ adenosine residues from oligoU tracts and leaves a 3′ cyclic phosphate that is not subjected to ring opening as with *S. cerevisiae* Usb1 (35,42). Human Usb1 was prepared exactly as described elsewhere (42). The conversion protocol involved resuspension of an RNA pellet from IDT in buffer E to a final concentration of 2.7 mM. A limiting amount of enzyme was used to ensure the product RNA was predominantly that of a single cleavage reaction. This was accomplished by adding 180 uL of RNA at 2.7 mM to 180 uL of human Usb1 at 32 uM in buffer F (100 mM NaCl, 10 mM HEPES acid, 10 mM sodium HEPES base, 40 % glycerol, 1 mM TCEP-HCl, 1 mM trisodium EDTA, pH ∼7) and incubation at 37 °C for 1 hour. The RNA was purified by 20 % polyacrylamide denaturing gel electrophoresis and anion exchange chromatography as above, with the exception that HiTrapQ wash buffer contained 100 mM NaCl, 10 mM HEPES acid and 10 mM sodium HEPES base, and the step elution buffer contained 1 M instead of 2 M NaCl. The eluted RNA was pooled and not subsequently adjusted prior to addition to Lsm2-8 (see below). The concentration of RNA was determined using an approximate molar extinction coefficient of 50,000 M^-1^cm^-1^ at 260 nm (41). All other RNAs used for co-crystallization experiments were purchased from IDT and resuspended in buffer E without further modification prior to addition to protein (see below).

The 5′-FAM labeled RNAs used for fluorescence polarization (*S. pombe* U6 nucleotides 91-100 and similar) were purchased from Integrated DNA Technologies and purified by urea PAGE and ion exchange as above for full-length U6, with the following exceptions: UV shadowing was not used (or required) to visualize the RNA after denaturing electrophoresis, concentration of RNA after anion-exchange chromatography was performed with 3 kDa MWCO spin filters (Amicon # UFC500308), and the final RNA concentration was estimated using UV absorption and an anticipated extinction coefficient of 75,000 M^-1^cm^-1^ at 495 nm (41). In the case of the cyclic phosphate probe, a limiting amount of human Usb1 was added to an adenosine terminated RNA as above.

### Fluorescence polarization binding assays

All fluorescence polarization binding assays were performed in buffer H (100 mM NaCl, 20 mM tris pH 8.2, 10 mM MgCl_2_, 1 mM TCEP HCl, 1 mM sodium azide, 0.1 mg/mL tRNA (Roche # 12172120), 0.1 mg/mL BSA (Ambion # AM2616) and 0.1 mg/mL sodium heparin (Sigma # N4784-250MG), pH ∼ 8) in black 96 well microplates (Greiner Bio-One # 655209) and imaged on a Tecan Infinite M1000Pro using an excitation wavelength of 470 nm and emission wavelength of 519 nm. For each sample, 100 uL of RNA at 2 nM was added to 100 uL of protein at a defined concentration between 0.4 nM and 5 uM. Fluorescence polarization was measured in duplicate from two independent titrations using different protein concentrations. Binding curves were fit using nonlinear regression in GraphPad Prism 4 to the following 4 parameter equation: FP = FP_min_ + (FP_max_ - FP_min_)/(1 + 10^((log*K*_d_ - log[protein])*H)), where FP_min_ and FP_max_ are the minimum and maximum polarizations, *K*_d_ is the binding dissociation constant, and H is the Hill coefficient. H was constrained to be 1 during non-linear regression. Depicted binding curves are normalized to FP_min_ and FP_max_. All raw binding data are provided in Supplementary Data File 1.

### Crystallization and structure determination of Lsm2-8 complexes

The Lsm2-8/RNA complexes were reconstituted by adding crude RNA from IDT to protein in an approximate two-fold stoichiometric excess. The complexes were then dialyzed overnight at 4 °C against 1 L of buffer I (50 mM NaCl, 10 mM MgCl_2_, 10 mM tris-HCl, 10 mM bis-tris base, 1 mM TCEP-HCl, pH ∼ 7) using 20 kDa MWCO membranes (Pierce # 66012). The dialyzed complexes were concentrated to approximately 10 mg/mL with 50 kDa MWCO spin filters (Amicon # UFC505008) prior to high-throughput crystallization screening on a Mosquito crystallization robot (TTP Labtech). Initial crystallization hits were obtained exclusively using the precipitant pentaerythritol propoxylate (5/4 PO/OH) in a MIDAS screen (Molecular Dimensions # MD1-59). Crystals were optimized by hanging drop vapor diffusion at 16 °C, using 2 μL of Lsm2-8/RNA complexes mixed with 2 μL of crystallization reagent containing 100-200 mM KCl, 50-100 mM HEPES pH 7.4, 35 % pentaerythritol propoxylate (5/4 PO/OH). Crystals were vitrified by direct immersion into liquid nitrogen.

Diffraction data were integrated using XDS (43). Space group determination was performed in *POINTLESS* (44). STARANISO (45) was used for merging and ellipsoidal truncation of the anisotropic diffraction data. *Phenix.xtriage* was used to assay potential twinning in the diffraction data (46). Initial phases were determined by molecular replacement using *Phaser* (47).

For the 3′ diol terminated structure (PDB 6PPN), three diffraction datasets were collected from two isomorphous crystals at 100 K on beamline 24-ID-E at the Advanced Photon Source. Molecular replacement was used to obtain initial phases with initial search templates PDB 4EMG (*S. pombe* Lsm3) (48), PDB 4EMH (*S. pombe* Lsm4) (48), PDB 4EMK (*S. pombe* Lsm5/6/7) (48), and homology models (49,50) of *S. pombe* Lsm2 and Lsm8 that were constructed from the corresponding *S. cerevisiae* orthologs (PDB 4C92 and 4M7D, respectively) (11,26). Molecular replacement used a single search model in which the above seven proteins were placed in a fixed orientation relative to one another to resemble the known architecture of the *S. cerevisiae* Lsm rings (11,25,26). Structure refinement was performed in *Phenix.refine* (46,51) using secondary structure restraints and TLS parameterization, with iterative rounds of manual model building in Coot (52,53) and additional automated refinement in *Phenix*.*refine*.

The 2′,3′-cyclic phosphate terminated structure (PDB 6PPP) was determined as above, but using addition of a different RNA prior to dialysis and using the above *S. pombe* Lsm2-8 structure for molecular replacement. Twelve diffraction datasets were collected from two isomorphous crystals at 100 K on beamline 21-ID-D at the Advanced Photon Source. Structure refinement was performed in *Phenix.refine* (46,51) using secondary structure restraints and TLS parameterization, in combination with reference model restraints to the best resolved Lsm2-8 ring in PDB 6PPN.

### Crystallization and structure determination of Lsm1-7 complexes

*S. pombe* Lsm1-7 complexes lacking the C-terminal 56 residues of Lsm1 were reconstituted with RNA as above, adding 5′-UUUUUA-3′ for PDB 6PPQ or 5′-AUUUUG-3′ for PDB 6PPV. Crystals were obtained by mixing 0.2 uL of complex with 0.2 uL of the following mixture: 20 mM sodium formate, 20 mM ammonium acetate, 20 mM trisodium citrate, 20 mM sodium potassium tartrate, 20 mM sodium oxamate, 100 mM sodium HEPES base, 100 mM MOPS acid, 10 % PEG 8,000 and 20 % ethylene glycol. Crystals were vitrified by direct immersion into liquid nitrogen.

The 3′ adenosine terminated structure (PDB 6PPQ) was determined by merging three diffraction datasets collected from a single crystal at 100 K on beamline 21-ID-D at the Advanced Photon Source. PDB 6PPN was used for molecular replacement. Structure refinement was performed in *Phenix.refine* (46,51) using secondary structure restraints. The 3′ adenosine binding pocket was first identified by residual Fo-Fc density after modeling and refining tetrauridylate into the four typical Sm-like pockets in Lsm1-7. Subsequent to incorporating the 3′ adenosine, the final electron density maps exhibited residual Fo-Fc density that could not be remediated by deletion of the adenosine or changing the identity of the adenosine to uridine. We therefore conclude the remaining Fo-Fc density is due to unmodeled dynamics in the 3′ adenosine binding pocket.

The 3′ guanosine terminated structure (PDB 6PPV) was determined from a single diffraction dataset collected from a single crystal at 100 K on beamline 21-ID-D at the Advanced Photon Source. PDB 6PPQ was used for molecular replacement and refinement was conducted as above. The 3′ guanosine binding pocket did not exhibit residual Fo-Fc density as above for adenosine.

For all structures presented here, simulated annealing omit maps were prepared to confirm the presence of bound RNA in the final models deposited into the Protein Data Bank. All figures were generated with PyMOL (http://www.pymol.org). Structural biology applications used in this project were compiled and configured by SBGrid (54). Electrostatic surface potentials were calculated using APBS (55) as implemented in PyMOL. All final coordinate sets and structure factors with calculated phases are provided in Supplementary Data File 2. A PyMol session with annotation matching that used throughout the manuscript is provided as Supplementary Data File 3.

## Results

### The Lsm2-8 complex specifically recognizes oligoU with a 2′,3′ cyclic phosphate

Lsm proteins were co-expressed in E. *coli* and orthogonal affinity tags were used to purify the Lsm1-7 and Lsm2-8 complexes (Supplementary Figure 1). For Lsm2-8, we demonstrated that the complex can bind to U6 snRNA and further associate with protein Prp24 to form the complete U6 snRNP (25). We wished to determine if the Lsm2-8 ring specifically recognizes the 2′,3′ cyclic phosphate group at the end of U6 snRNA, and if so, how. We therefore compared the binding affinities of Lsm2-8 for oligoribonucleotides corresponding to the 3′ oligoU tail of U6 terminating with a 2′,3′ cyclic phosphate group, a 3′ phosphate, or an unmodified 3′ hydroxyl (Figure 1b, Supplementary Table 1).

We find that Lsm2-8 binds tightly to an RNA oligonucleotide with a 2′,3′ cyclic phosphate group, and 4-fold less tightly to the unmodified RNA with a 3′ hydroxyl (*K*_d_ = 26 and 100 nM, respectively) (Figure 1b). This 4-fold difference suggests specific recognition of the 2′,3′ cyclic phosphate group. Interestingly, Lsm2-8 binds to an oligonucleotide terminating in a 3′ phosphate least tightly of all with a *K*_d_ of 152 nM. Thus, the Lsm2-8 complex preferentially binds to oligonucleotides terminating in a 2′,3′ cyclic phosphate group while still retaining ability to bind to unmodified RNA, and can somehow strongly discriminate between 2′,3′ cyclic phosphate and non-cyclic 3′ phosphate groups.

### Structures of Lsm2-8-RNA complexes

We crystallized the Lsm2-8 complex bound to UUUUU-3′-OH and UUUUU>p and determined their structures by x-ray diffraction to resolutions of 1.9 and 2.3 Å, respectively (Table 1). In the Lsm2-8 complex bound to the UUUUU-3′-OH RNA, the 5′ uridine is bound by Lsm4, and the next three uridines are bound sequentially by Lsm8, Lsm2, and Lsm3 (Figure 2a,c). Uridine binding involves stacking and an extensive hydrogen bonding network as previously described (25,26) (Figure 2e). The last uridine with the 3′-OH group is disordered except for its 5′ phosphate as evidenced by weak electron density for both the ribose and uracil groups (Supplementary Figure 2a). We confirmed that the RNA remains completely intact after prolonged incubation with Lsm2-8, further supporting the idea that the last uridine is covalently attached but disordered in the structure (Supplementary Figure 2b).

**Table 1.** Diffraction data collection and structure refinement statistics

| Protein<br>RNA | Lsm2-8<br>UUUUUU | Lsm2-8<br>UUUUU>p | Lsm1<sub>Δ56C</sub>-7<br>UUUUUA | Lsm1<sub>Δ56C</sub>-7<br>AUUUUG |
| --- | --- | --- | --- | --- |
| PDB ID | 6PPN | 6PPP | 6PPQ | 6PPV |
| Wavelength (Å) | 0.9792 | 1.0781 | 0.9762 | 0.9763 |
| Resolution range (Å) | 168-1.90 (2.08-1.90) | 78.0-2.33 (2.73-2.33) | 98.8-1.81 (1.88-1.81) | 98.7-2.05 (2.11-2.05) |
| Space group | *P*2<sub>1</sub>2<sub>1</sub>2 | *P*2<sub>1</sub>2<sub>1</sub>2 | *P*3<sub>1</sub>21 | *P*3<sub>1</sub>21 |
| Unit cell (Å) | 70.3 135.4 168.0 | 70.3 139.7 153.9 | 68.9 68.9 296.3 | 69.0 69.0 296.1 |
| Total reflections* | 1,101,742 (46,024) | 2,493,903 (104,496) | 2,384,574 (75,834) | 1,117,285 (89,763) |
| Unique reflections* | 80,785 (4,039) | 28,712 (1,433) | 75,844 (4,347) | 52,874 (4,047) |
| Multiplicity* | 13.6 (11.4) | 86.9 (72.9) | 31.4 (17.4) | 21.1 (22.2) |
| Completeness (%)* |  |  |  |  |
| spherical | 63.3 (13.4) | 43.7 (5.9) | 100.0 (99.4) | 100 (100) |
| ellipsoidal | 96.2 (74.4) | 94.9 (74.6) | NA | NA |
| Anisotropic $\Delta B$ (Å<sup>2</sup>) | 32.4 | 83.4 | 10.1 | 7.4 |
| Mean $I/\sigma(I)$* | 17.2 (1.8) | 16.0 (2.2) | 15.4 (0.4) | 11.6 (0.9) |
| $R_{\text{merge}}$* | 0.074 (1.355) | 0.293 (3.347) | 0.125 (7.075) | 0.159 (4.145) |
| $R_{\text{pim}}$* | 0.021 (0.401) | 0.032 (0.392) | 0.022 (1.698) | 0.036 (0.897) |
| $CC_{1/2}$* | 0.999 (0.729) | 0.999 (0.876) | 0.999 (0.409) | 0.999 (0.380) |
| **Refinement** |  |  |  |  |
| Resolution | 67.7-1.91 (1.98-1.91) | 51.7-2.33 (2.41-2.33) | 34.5-1.81 (1.88-1.81) | 42.1-2.05 (2.12-2.05) |
| $R_{\text{work}}/R_{\text{free}}$* | 0.20/0.23 (0.34/0.27) | 0.22/0.28 (0.62/0.63) | 0.19/0.22 (0.36/0.40) | 0.21/0.25 (0.35/0.38) |
| Total atoms | 8,877 | 8,841 | 4,677 | 4577 |
| macromolecules | 8,783 | 8,815 | 4,375 | 4387 |
| ligands | 0 | 0 | 0 | 0 |
| water | 94 | 26 | 302 | 190 |
| RMS(bonds) | 0.009 | 0.013 | 0.006 | 0.005 |
| RMS(angles) | 1.19 | 1.95 | 0.89 | 0.87 |
| Ramachandran favored | 97.9 % | 98.0 % | 98.3 % | 96.3 |
| Ramachandran outliers | 0 % | 0 % | 0 % | 0 % |
| Average ADP (Å<sup>2</sup>) | 66.9 | 85.7 | 51.5 | 50.9 |
| macromolecules | 67.1 | 85.8 | 50.9 | 50.8 |
| ligands/ions | NA | NA | NA | NA |
| solvent | 47.5 | 46.0 | 60.6 | 55.4 |

STARANISO (71) was used to perform ellipsoidal truncation for datasets 6PPN and 6PPP. After ellipsoidal truncation the weakest direction of diffraction data extended to 2.75 Å for PDB 6PPN and 3.60 Å for PDB 6PPP. Data from ellipsoidal truncation accompany the coordinate sets deposited in the Protein Data Bank. For PDBs 6PPN and 6PPP, merged diffraction data that has not been subjected to ellipsoidal truncation is available as Supplementary Data File 2, along
with the final refined coordinates, structure factors and phases for all structures reported here. A summary PyMOL session file is available as Supplementary Data File 3.

**Figure 2.**
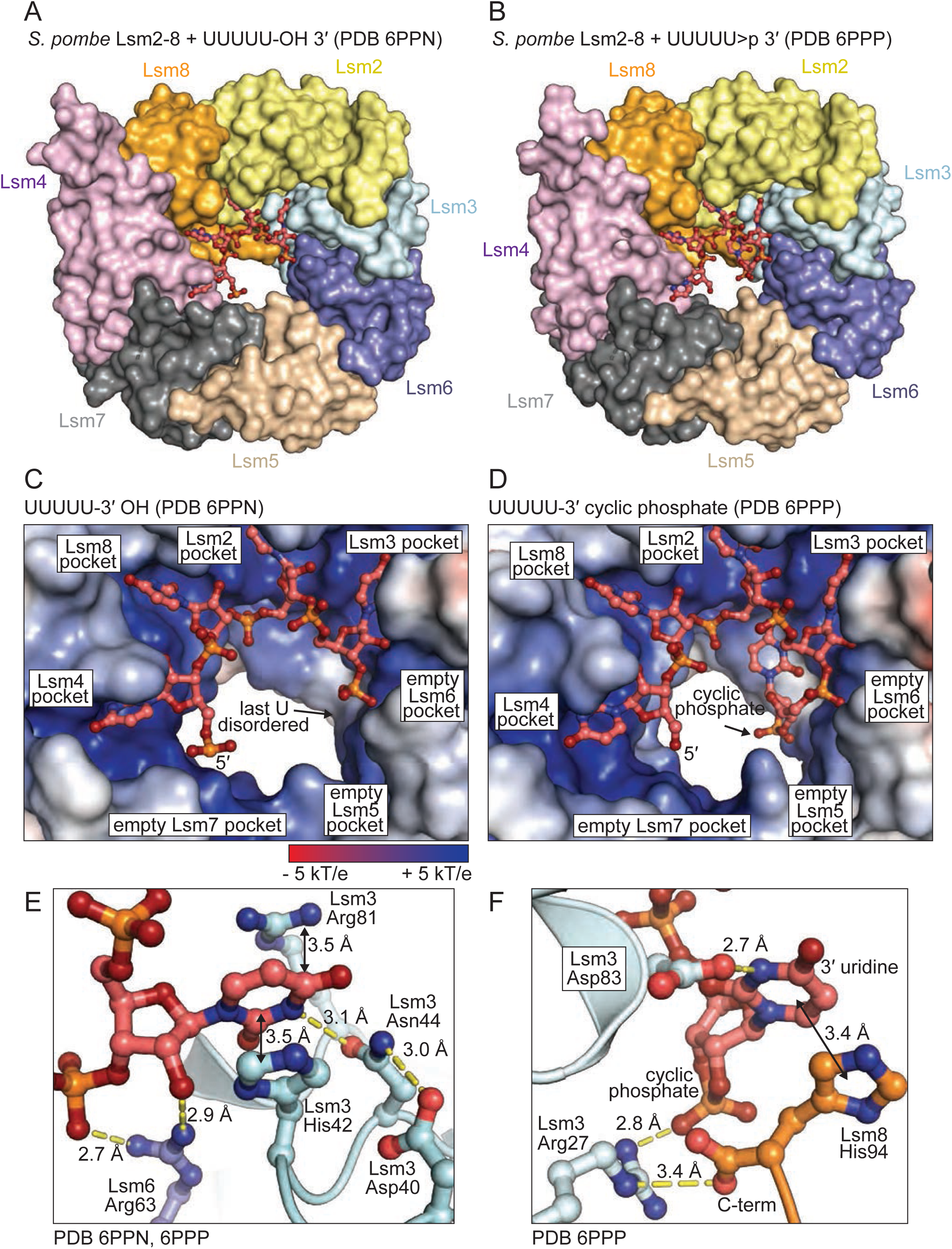
Structures of *S. pombe* Lsm2-8 bound to pentauridylate with a 2’,3’ OH or 2’,3’ cyclic phosphate. A) Overview of Lsm2-8 bound to “unprocessed” U6 snRNA 3’ terminus. B) Overview of Lsm2-8 bound to “mature” U6 snRNA 3’ terminus. C) Detail of Lsm2-8 interface with “unprocessed” RNA. The 3’ terminal uridine is disordered and thus not visible in the final electron density maps. D) In contrast, the mature U6 3’ end, with a 2’,3′ cyclic phosphate, shows electron density for the terminal nucleotide. E) The Sm-like pocket in Lsm3 binds RNA as observed previously in other Lsm2-8 complexes from *S. cerevisiae*. F) In contrast, the 3′ uridine cyclic phosphate has a unique binding mechanism relative to the other four uridines, including a stacking interaction with the C-terminal histidine of Lsm8.

In contrast to the Lsm2-8 complex with unmodified RNA, the structure of the Lsm2-8 ring bound to UUUUU>p reveals that the last nucleotide is highly ordered (Supplementary Figures 2c,d). The cyclic phosphate causes the RNA chain to make a sharp turn of nearly 180°, which positions the terminal nucleobase to stack over the last histidine at the C-terminus of Lsm8. The terminal uracil base adopts an unusual *syn* conformation and forms a hydrogen bond to the Lsm3-Asp83 side chain (Figure 2f). Lsm3-Arg27 forms a hydrogen bond to a non-bridging oxygen on the terminal cyclic phosphate group and makes a salt bridge with the Lsm8 C-terminal histidine carboxyl group. This binding mechanism is markedly different from that in the *S. cervisiae* Lsm2-8 complex with a non-cyclic phosphate, where the terminal uridine adopts an *anti* conformation and lacks direct contacts with the corresponding arginine and aspartate residues (25) (Supplementary Figure 3).

### RNA binding properties of the Lsm1-7 complex

Lsm1-7 shares 6 out of 7 subunits with Lsm2-8 and has all but one of the uridine binding pockets present in Lsm2-8. The *S. cerevisiae* Lsm1-7 structure in the absence of RNA has been determined and displays an overall structure that is very similar to Lsm2-8 (Figure 3). We therefore reasoned that Lsm1-7 and Lsm2-8 might bind to similar RNA sequences. However, when superimposing the structures of *S. cerevisiae* Lsm2-8 bound to RNA and Lsm1-7, there is steric clash between the 3′-end of the RNA and the C-terminal domain of Lsm1 (Figure 3b). This may explain why the *S. cerevisiae* Lsm1-7 ring alone has been reported to not bind to RNA (24).

**Figure 3.**
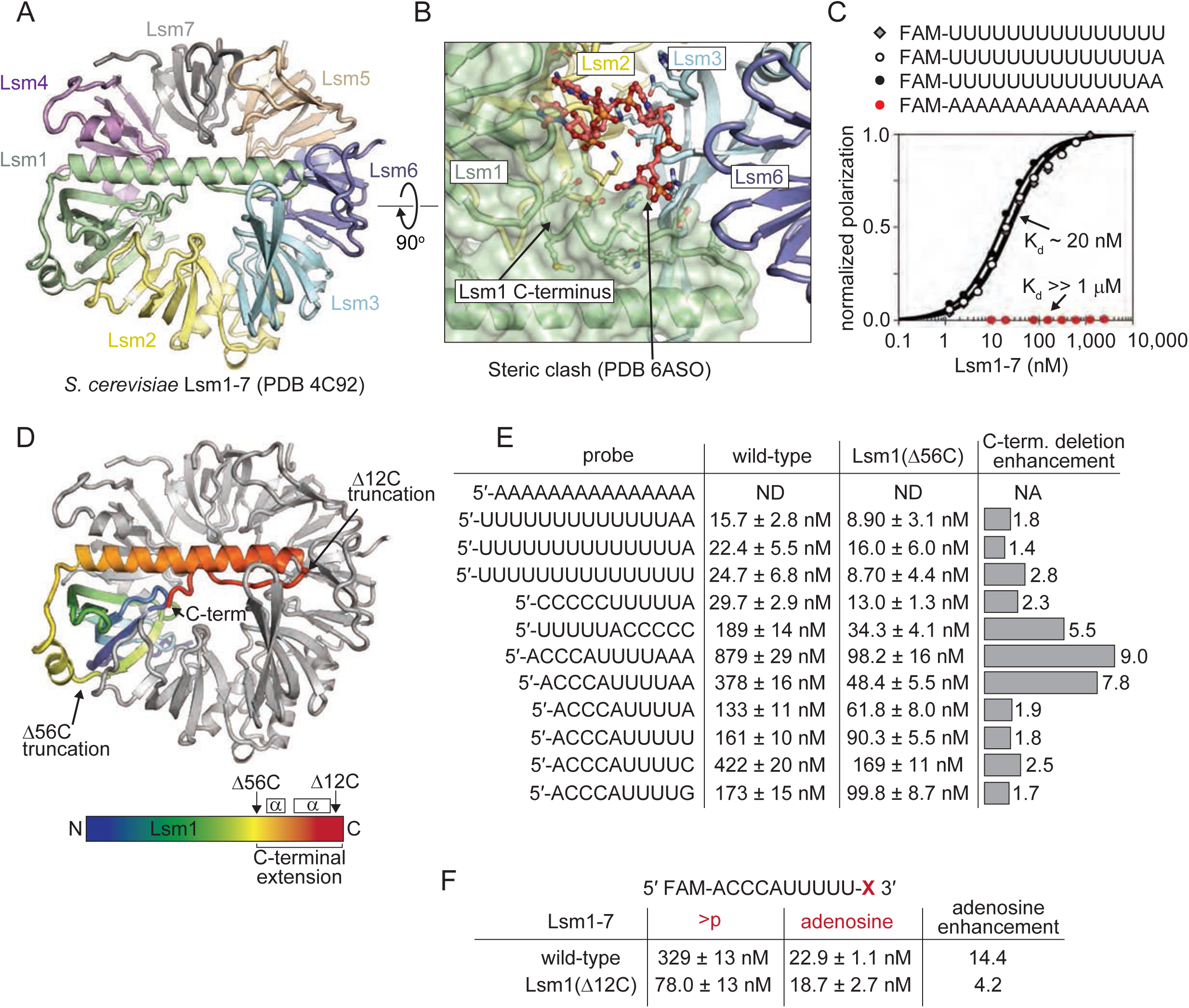
The elongated C-terminal region of Lsm1 in the *S. pombe* Lsm1-7 ring attenuates RNA binding. A) Structure of the *S. cerevisiae* Lsm1-7 ring in the absence of bound RNA (11). The C-terminal region of Lsm1 crosses the distal face of the ring, occluding the central pore. B) Superposition of U6 snRNA from the homologous *S. cerevisiae* Lsm2-8 ring with the *S. cerevisiae* Lsm1-7 ring. The alignment in this figure was achieved by superposition of Lsm2, Lsm3 and Lsm6 components of the rings, which share a pairwise r.m.s.d of 0.5 Å. A steric clash is visible between the 3′ uridine phosphate and the C-terminal region of Lsm1. C) *In vitro* fluorescence polarization binding assays show that *S. pombe* Lsm1-7 tightly binds to polyuridylate tracts without accessory proteins, like Pat1. In contrast, *S. pombe* Lsm1-7 lacks detectible affinity for polyadenylate. D) Designed truncations of the Lsm1 C-terminus. Lsm1 is colored blue to red from the N and C termini, respectively, and regions selected for truncation are annotated in the figure, resulting in truncation of the last 12 residues that fold back into the pore, or truncation of the entire helical region that spans the distal face of the ring. E) Binding assays showing that deletion of the Lsm1 C-terminal region generally enhances binding affinity for RNAs that harbor polyuridine tracts, and the relative enhancement is greatest for the weakest binding RNAs. F) A strong preference for an adenosine 3’ terminus over a uridine cyclic phosphate 3’ terminus is diminished upon deletion of the C-terminal 12 residues of Lsm1.

Since *S. pombe* Lsm1-7 has been observed to bind to oligoU RNAs as well as oligoA RNAs in the presence of Pat1 (12), we first analyzed binding to four 15mers: U15, U14A, U13AA, and A15 under stringent binding conditions including 0.1 mg/mL competitor tRNA, 0.1 mg/mL sodium heparin and 0.1 mg/mL BSA. We find that U15, U14A and U13AA all bind tightly to Lsm1-7; in contrast, A15 does not bind in our assay (Figure 3c). Owing to the observed potential for steric clash between RNA and the C-terminus of Lsm1, we also compared RNA binding to a variant of Lsm1-7 in which the last 56 amino acids of Lsm1 were removed (Lsm1_Δ56C_-7) (Figure 3d). Lsm1_Δ56C_-7 binds even more tightly to the oligoU RNAs and, like Lsm1-7, also does not bind to A15 (Figure 3e). Upon observing high affinity binding of Lsm1-7 to oligoU RNAs, we assayed RNAs with shorter U-tracts for binding. Lsm1-7 and Lsm1_Δ56C_-7 binds tightly to the oligo CCCCCUUUUUA (*K*_d_ = 29.7 and 13 nM, respectively), which is comparable to the binding observed for longer oligoU tracts (Supplementary Table 1). The oligo ACCCAUUUUU binds less tightly than U15, indicating that the length of the oligoU tract plays a role in binding affinity.

In general, the Lsm1_Δ56C_-7 construct has higher affinity for RNAs than Lsm1-7 (Figure 3e). When the oligoU tract is at or near the 3′ end, the difference in binding affinities between Lsm1-7 and Lsm1_Δ56C_-7 is approximately 2-fold. However, if the oligoU tract is followed by two or more 3′ nucleotides, the binding enhancement afforded by the Lsm1 C-terminal deletion becomes much more significant (5 to 9-fold). For example, the oligonucleotide UUUUUACCCCC binds poorly to Lsm1-7 (*K*_d_ = 189 nM) and tightly to Lsm1_Δ56C_-7 (*K*_d_ = 34 nM) (Figure 3e). These data indicate that high affinity binding sites for the Lsm1-7 complex must be at the 3′ termini of RNA. On the other hand, the C-terminally truncated Lsm1-7(Lsm1_Δ56C_-7) complex can bind tightly to the UUUUA sequence irrespective of its position in the RNA. Thus, the Lsm1 C-terminal extension functions to prevent high affinity binding to RNA, except for 3′ terminal sites. We further tested the effect of 3′-end nucleotide identity on binding and found that tetrauridylate followed by a single adenosine binds more tightly than an RNA with a pentauridylate tract (Figure 3e).

Interestingly, we observe that the C-terminal 12 residues of Lsm1 strongly discriminates against cyclic phosphate RNAs. Comparison of a monoadenylated RNA and a cyclic phosphate RNA shows an approximate 14-fold reduction in binding affinity for the cyclic phosphate RNA, which is substantially attenuated by deleting only a small region from the C-terminus of Lsm1 (Figures 3d,f). We conclude that the C-terminal 12 amino acids of Lsm1 are important for the binding specificity of Lsm1-7.

### Structures of Lsm1-7 bound to RNA

We were unable to obtain crystals of wild-type Lsm1-7 in complex with RNA. Since deletion of the C-terminus of Lsm1 generally enhances binding affinity (Figure 3e), we therefore crystallized the Lsm1_Δ56C_-7 variant of the Lsm1-7 complex bound to the U-tract RNAs UUUUUA and AUUUUG and determined their structures to resolutions of 1.8 and 2.1 Å, respectively (Figure 4 and Table 1). In the structure of UUUUUA bound to Lsm1_Δ56C_-7, the first uridine is disordered and not visible in the electron density (Figure 4a,c; Supplementary Figure 4). The next four uridines occupy pockets in Lsm4, Lsm1, Lsm2 and Lsm3, respectively. The terminal adenosine reaches across Lsm6 to form a hydrogen bond to Asn66 of Lsm5 (Figure 4e). This is the first observation of Lsm5 interacting with RNA; Lsm5 is not utilized in the Lsm2-8 structures. Furthermore, Lsm5 is the only Lsm protein out of the 8 studied here that has a non-canonical nucleobase binding pocket because it is missing the arginine that typically forms a cation-pi stack with uracil (14,25,26) (Figure 2e). In place of arginine, Lsm5 has an asparagine (Asn66) that forms a hydrogen bond to the terminal adenine. Interestingly, this non-canonical asparagine in the Lsm5 binding pocket is highly conserved (Supplementary Data File 4).

**Figure 4.**
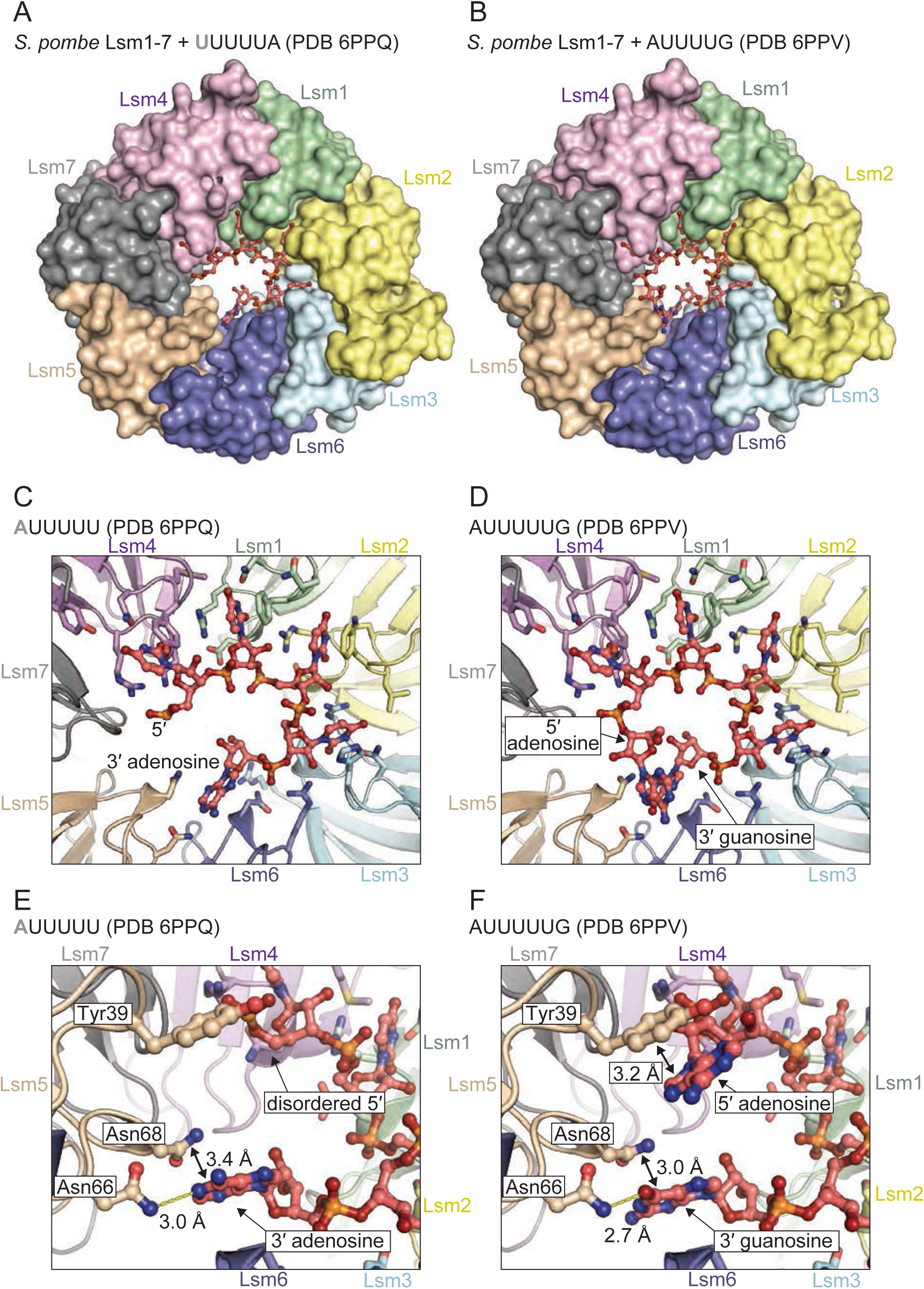
The structure of *S. pombe* Lsm1-7 bound to RNA explains the preference for a 3’ terminal purine. A) Overview of Lsm1-7 bound to UUUUA. B) Overview of Lsm1-7 bound to AUUUUG. C) RNA-binding interface of Lsm1-7 bound to UUUUA, showing that four uracil bases bind in the same manner as in the Lsm2-8 complex, while the adenosine binds into non-Sm pocket in Lsm5. D) Lsm1-7 bound to AUUUUG, showing a similar binding mechanism as in panel C. E,F) Detailed view of the non-Sm Lsm5 binding pocket occupied by adenine or guanine. In both cases, the 3′ purine is coordinated through hydrogen bonding with Lsm5-Asn66 and stacking with Lsm5-Asn68.

In the structure of Lsm1_Δ56C_-7 bound to AUUUUG, all nucleotides are visible within the electron density map. The path of the RNA around the interior of the Lsm1-7 torus is a right-handed helix, with the phosphodiester backbone in the center and the nucleobases splayed out into binding pockets. The six nucleotides of the RNA make an almost complete turn with the last nucleotide 6 Å below the first (Figures 4d,f). The 5′ adenosine interacts with the non-canonical binding pocket on Lsm5 by stacking on Tyr39 (Figure 4f). The 3′ G also interacts with Lsm5 by forming a hydrogen bond to Asn66 in a manner that is similar to the interaction observed for the terminal adenine of UUUUUA, except the asparagine side chain donates a hydrogen bond to the guanine oxygen instead of an adenine nitrogen (Figures 4e,f). The structures explain why oligonucleotides harboring these sequences bind with nearly identical affinities to Lsm1_Δ56C_-7 (Figure 3e).

Since the terminal purines are both within hydrogen bonding distance of Lsm5 Asn66 and are in van der Waals contact with Lsm5 Asn68 (Figures 4e,f), we tested the contribution of these interactions to binding by mutating Lsm5 Asn66 and Asn68 to alanine. The double mutant has a small but measurable effect upon binding RNA (*K*_d_ = 23 vs 41 nM) (Supplementary Table 1). We also tested the binding specificity of the double mutant by comparing its ability to bind an RNA terminating with a cyclic phosphate vs the terminal adenosine. While Lsm1-7 can discriminate between these RNAs with a 14-fold difference in affinity, the double mutant shows only a 7-fold difference, corresponding to a 2-fold loss in binding specificity (Supplementary Table 1).

### Lsm1-7 loads onto RNA from 3′ ends and is blocked by secondary structure

In the crystal structures, the 5′ end of the RNA hexamer is nearest the “proximal face” of Lsm1-7 while the 3′ end is nearest the “distal face” (Figure 5a), the same orientation as in the Sm ring where the bound RNA passes completely through the center of the torus. This similarity suggests that single stranded RNA may be able to pass through the Lsm1-7 torus. Moreover, we reasoned that binding may be affected by adjacent secondary structure, particularly if the complex were to load onto RNA in a directional manner. We therefore created RNA constructs with UUUUA binding sites and stem-loop structures at either the 5′ or 3′ termini or both (Figure 5b). We find that both Lsm1-7 and Lsm1_Δ56C_-7 can only bind tightly to the RNA with a 5′ stem-loop and single stranded 3′ end (*K*_d_ = 70 and 32 nM, respectively) (Figure 5c). When the 5′ end is single stranded with a hairpin at the 3′ end, binding is severely weakened by 25-fold or more (*K*_d_ > 1 µM). This loss in binding affinity cannot be attributed to the 5′ cytidines, which do not affect binding to single stranded RNA (Figure 3e). With hairpins at both ends, the binding is also severely weakened. We therefore conclude that both Lsm1-7 and Lsm1_Δ56C_-7 load directionally onto the 3′ ends of single-stranded RNAs, and that binding is effectively blocked by downstream secondary structure. Comparison of surface electrostatics in Lsm1_Δ56C_-7 shows the proximal face has more electropositive surface area than the distal face of the ring (Supplementary Figure 5a), which may facilitate initial contact with RNA. Furthermore, comparison to the human Sm ring bound to U4 snRNA (56,57) shows how directional threading of the RNA through the ring may be facilitated by conserved distal face contacts, as several residues in SmD1 and Sm2 that interact with RNA are also present in Lsm rings (Supplementary Figure 5b).

**Figure 5.**
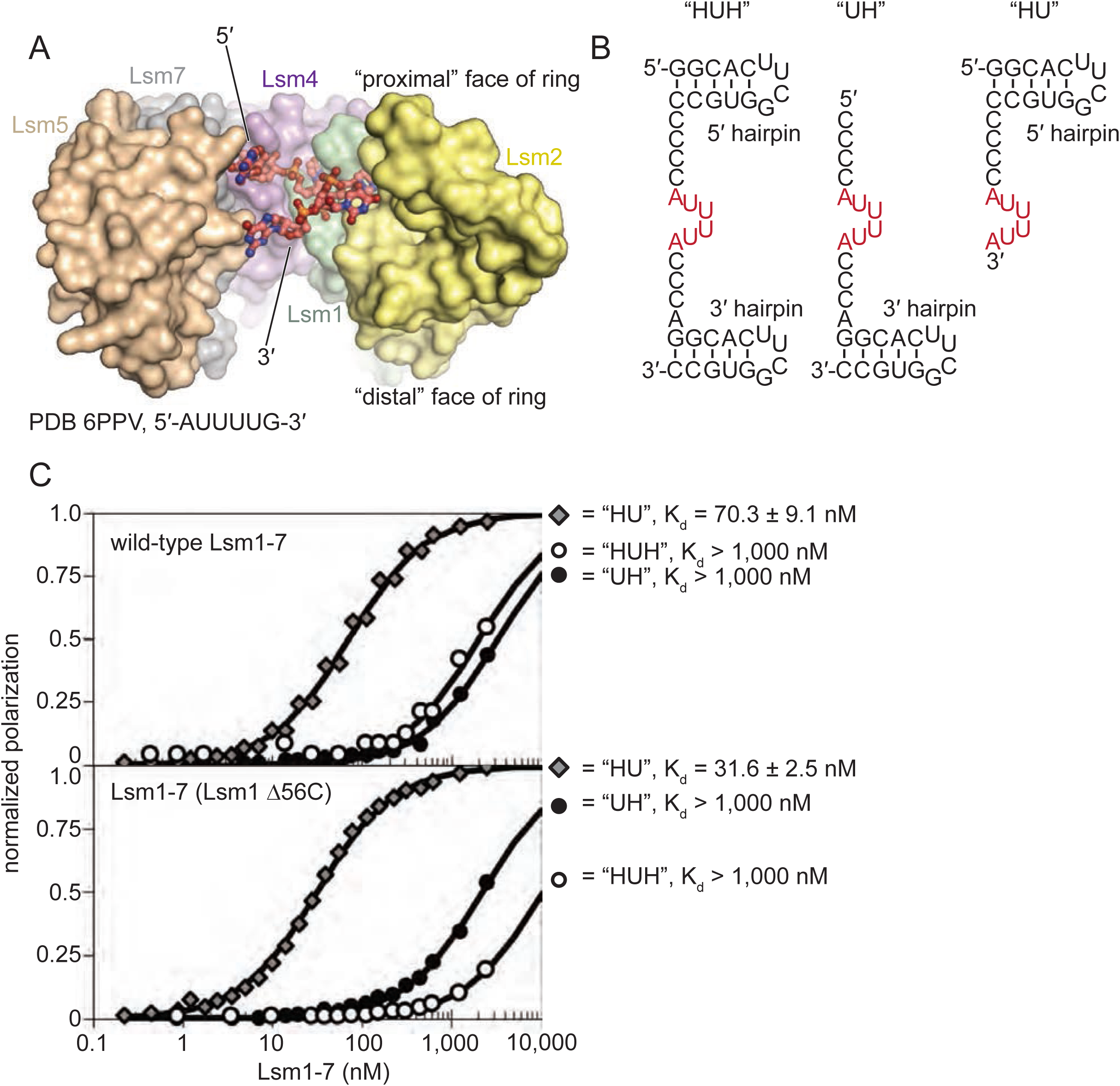
Lsm1-7 loads single stranded RNA from the 3′ end. A) Cut-away view of the pore of Lsm1-7 bound to AUUUUG. Lsm3 and Lsm6 are omitted for clarity. B) Synthetic RNAs used for binding tests, in which the 3′ or 5′ end is blocked from binding (or threading through) the pore of the Lsm1-7 ring by the presence of the hairpin structure. C) Wild-type Lsm1-7 tightly binds the 5′ hairpin RNA, and has weak binding for RNAs harboring a 3′ hairpin. Deletion of the C-terminal region of Lsm1 does not significantly alter the binding specificity with respect to 3′ or 5′ hairpins.

## Discussion

### Structural basis for RNA recognition by Lsm2-8

Phylogenetic analyses suggest that the Lsm/Sm family of proteins arose approximately 2.5 billion years ago through two waves of gene duplication, the first wave resulting in the Lsm genes from which the Sm proteins subsequently diversified (58). As Lsm and Sm proteins interactions are common to all spliceosomal RNAs, it is likely that these interactions arose early during the evolution of the spliceosome. The *S. pombe* Lsm2-8 complex is highly similar to human and other organisms (Supplementary Data File 4), so we expect the structural and functional data presented here to be broadly general in eukaryotic biology.

The molecular basis for Lsm2-8 binding is rationalized by the unique structural features observed in the Lsm2-8 complex bound to UUUUU>p. The 2′,3′ cyclic phosphate group bends the RNA chain around and stabilizes an unusual *syn* conformation of the terminal uracil base. Quantum chemical calculations of 2′,3′ cyclic UMP indicate that the cyclic phosphate stabilizes the *syn* conformation of uridine to be significantly more favored over the *anti* conformation (59). This *syn* conformation of the terminal uridine facilitates stacking on the Lsm8 C-terminal histidine and allows for a hydrogen bond to Lsm3 Asp83, which could not happen in an *anti* conformation. Consistent with its central role in directly binding to the cyclic phosphate, Lsm3 Arg27 is 100% conserved (25). The C-terminal histidine residue of Lsm8 that stacks with the terminal *syn* uracil nucleobase is also strongly conserved in almost all eukaryotes (Supplementary Data File 4). It is therefore clear that the C-terminus of Lsm8 and nearby residues are important for specifically recognizing post-transcriptionally modified U6 snRNA (25,35).

Although *S. pombe* Lsm2-8 binds most tightly to oligoU with a 2′,3′ cyclic phosphate, it also must recognize heterogenous RNA sequences and different 3′-ends *in vivo*. For example, Lsm2-8 binds to telomerase RNA (TER1), which has a stretch of 3-6 uridines ending with a 3′-OH. Consistent with this, Lsm2-8 still binds to oligoU RNA ending in 3′-OH, with a *K*_d_ of 100 nM. We find that Lsm2-8 also binds tightly to 3′ monoadenylated RNA (Supplementary Table 1). Interestingly, we note that 3’ monoadenylated U6 is present in human cells and is the preferred substrate for Usb1 (42,60,61).

### Unique RNA binding properties of the Lsm1-7 complex

In contrast to Lsm2-8, the Lsm1-7 complex strongly discriminates against RNAs with a 2′,3′ cyclic phosphate by 14-fold (Supplementary Table 1 and Figure 3f). This may be to avoid certain RNAs in the cytoplasm; for example, tRNA splicing proceeds through a cyclic phosphate intermediate (62). Our binding measurements indicate that Lsm1-7 binds tightly to RNAs containing UUUUR and loads onto RNA from the 3′ end. This is consistent with previous reports of Lsm1-7 binding to UA rich regions of viral genomes to regulate translation (22,63). Lsm1-7 binds tightly to the sequence AUUUUR (Figure 3e), and this sequence is highly reminiscent of sequences found in 3′ UTR regions that undergo adenine/uridine-rich element (ARE) mRNA decay (64,65). Consistent with this, knock-down of Lsm1 inhibits ARE-mediated decay (66) and depletion of the Lsm1-7 binding protein Pat1b upregulates mRNAs with AREs (67).

Lsm1-7 preferentially loads onto single-stranded RNA from the 3′ end. The trajectory of the RNA in Lsm1_Δ56C_-7 suggests that the RNA can thread completely through the ring, provided the Lsm1 C-terminal helix is removed or displaced. Indeed, our binding data show that RNAs with high affinity binding sites followed by additional 3′ nucleotides (e.g., AAA or CCCCC) bind well to Lsm1_Δ56C_-7 but not Lsm1-7. This is likely due to steric clash involving the Lsm1 C-terminal region (Figure 3b). The Lsm1 C-terminal region may act as a gate to allow threading of the RNA through the ring, followed by scanning via a one-dimensional search process, or sliding (68,69). Accordingly, there are conserved residues in the Lsm and Sm rings, the latter of which have previously been observed to interact with “threaded” single stranded RNA (Supplementary Figure 5).

### A model for Pat1 stimulation of RNA binding by Lsm1-7

Taken together, our data suggest an allosteric model for Pat1-mediated stimulation of Lsm1-7 binding (12,24). In this model, Pat1 binding displaces the Lsm1 C-terminal domain to stimulate RNA binding affinity and relax specificity (Figure 6). This model is consistent with existing structural data indicating that the C-terminal domain of Pat1 binds to Lsm2 and Lsm3 (11,15), close to where the Lsm1 C-terminal helix reaches across to dock on the other side of the ring (Figure 6a). This positioning would place the middle domain of Pat1, which is not present in the structure but is required for high affinity binding to oligoA RNA (12), very close to the C-terminal helix of Lsm1. Our model is also consistent with the observation that mutation of the Pat1 interaction surface with Lsm2-Lsm3 destroys the ability of Pat1 to stimulate the RNA-binding activity of *S. pombe* Lsm1-7 (12) and leads to defects in mRNA degradation in *S. cerevisiae* (11,15). In *S. cerevisiae*, the C-terminal alpha-helical extension of Lsm1 is required for high affinity binding to oligoA RNA in the presence of Pat1 (70).

**Figure 6.**
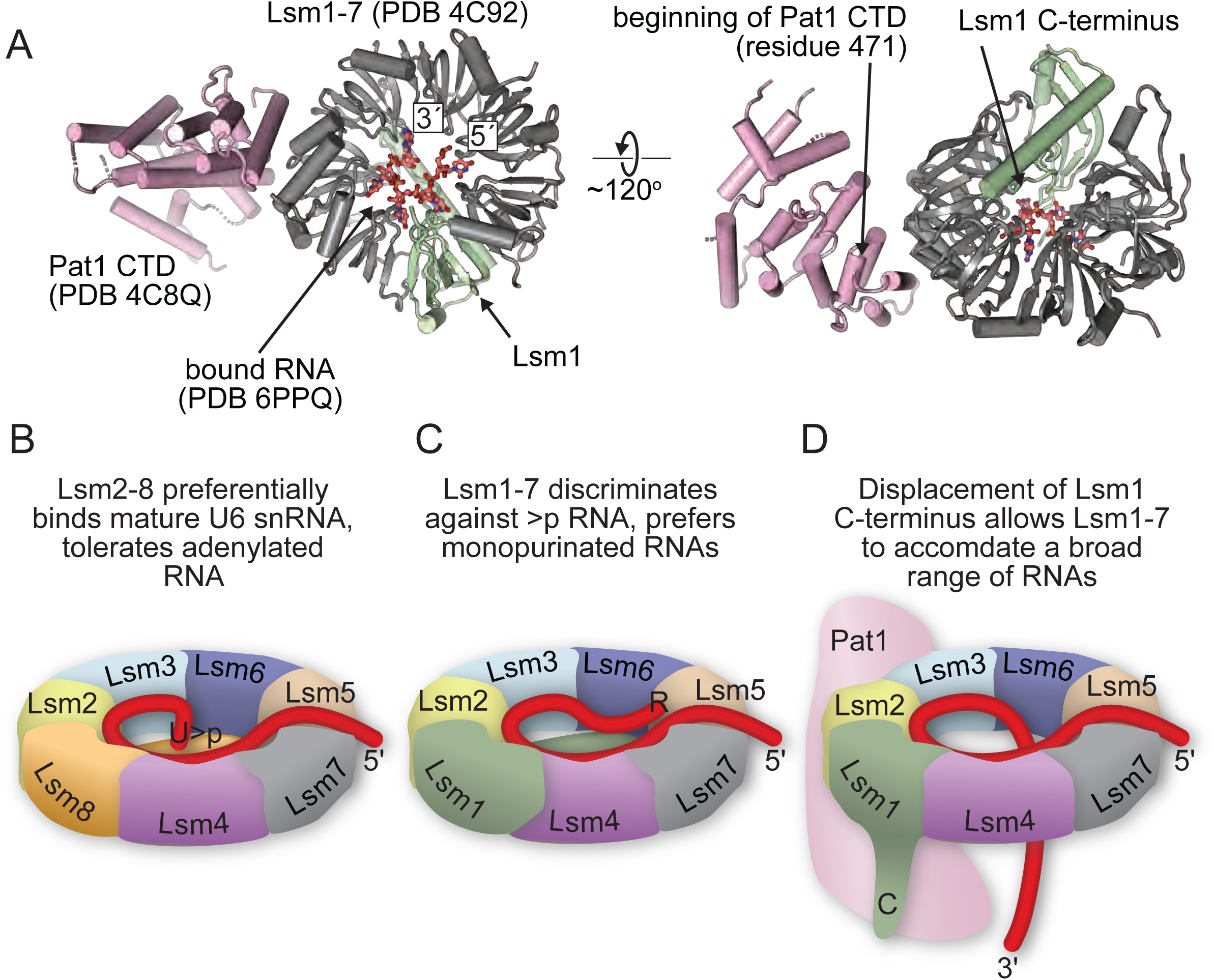
Proposed model for Pat1-regulated gating of the Lsm1-7 RNA-binding pore. A) Relative orientations of RNA, Lsm1-7 and Pat1 proteins, derived from structures of *S. pombe* Lsm1-7 with RNA and *S. cerevisiae* Lsm1-7 with the C-terminal domain of Pat1 (11). B) Model for RNA binding specificity in Lsm rings. The Lsm2-8 ring contains a uridine-cyclic phosphate binding site that includes the C-terminus of Lsm8 and endows specificity for Usb1 processed U6 snRNA. C) In contrast, the C-terminal region of Lsm1 in the Lsm1-7 ring antagonizes association of the ring with uridine-cyclic phosphate terminated RNA, while allowing association of the ring with 3′ monopurinated polyuridylate tracts. D) Deletion of the C-terminal region of Lsm1 allows the ring to associate with a broader range of RNAs. *In vivo*, the role of Lsm1 truncation may be mimicked by displacement of the C-terminus by association of the ring with the Pat1 co-factor. It remains to be seen if RNA is capable of threading all the way through the pore of the RNA, as depicted here, or rather enters and exits the pore on the proximal face alone.

In summary, the Lsm proteins are essential RNA binding proteins that initiate the formation of molecular assemblies involved in major pathways of gene expression, including pre-mRNA splicing and mRNA decay. Understanding how these proteins recognize their RNA targets is therefore an important aspect of eukaryotic biology. We quantitatively demonstrate that the Lsm1-7 and Lsm2-8 complexes achieve strikingly different functional properties, despite similar quaternary structures and sharing of subunits. By elucidating the structures of these complexes bound to RNA, we establish the molecular basis for RNA-protein interactions that are fundamental to eukaryotic gene expression.

## Acknowledgements

We are grateful to Aaron Hoskins, members of the Brow and Butcher laboratories, Elsebet Lund and Stephen Floor for helpful discussions and critical reading of the manuscript. Use of the Advanced Photon Source, an Office of Science User Facility operated for the U.S. Department of Energy (DOE) Office of Science by Argonne National Laboratory, was supported by the U.S. DOE under Contract No. DE-AC02-06CH11357. Use of NE-CAT was supported by the National Institutes of Health (NIH) grants P41 GM103403 and S10 RR029205. Use of LS-CAT was supported by NIH grant 085P1000817. We acknowledge the use of the SAXS Core facility of Center for Cancer Research (CCR), National Cancer Institute (NCI). This project has been funded in whole or in part with federal funds from the National Cancer Institute, National Institutes of Health, under contract HHSN26120080001E. This research used 12-ID-B beamline of the Advanced Photon Source, a U.S. Department of Energy (DOE) Office of Science User Facility operated for the DOE Office of Science by Argonne National Laboratory under Contract No. DE-AC02-06CH11357. Fluorescence polarization data were obtained at the University of Wisconsin-Madison Biophysics Instrumentation Facility, which was established with support from the University of Wisconsin-Madison and grants BIR-9512577 (NSF) and S10RR13790 (NIH). This study was supported by NIH grants R35 GM118075 to D.A.B. and R35 GM118131 to S.E.B.

## Author Contributions

E.J.M., J.M.V, S.M.H., Y.N. prepared reagents. E.J.M. collected diffraction data and determined the crystal structures. E.J.M and J.M.V performed *in vitro* binding assays. E.J.M. and S.E.B. supervised the work. E.J.M., D.A.B and S.E.B. wrote the manuscript.

## Author Information

Other data supporting the findings of this manuscript are available from the corresponding authors upon request. The authors declare no competing financial interests. Correspondence and requests for materials should be addressed to E.J.M and S.E.B.

## Figure captions

**Supplementary Table 1.**
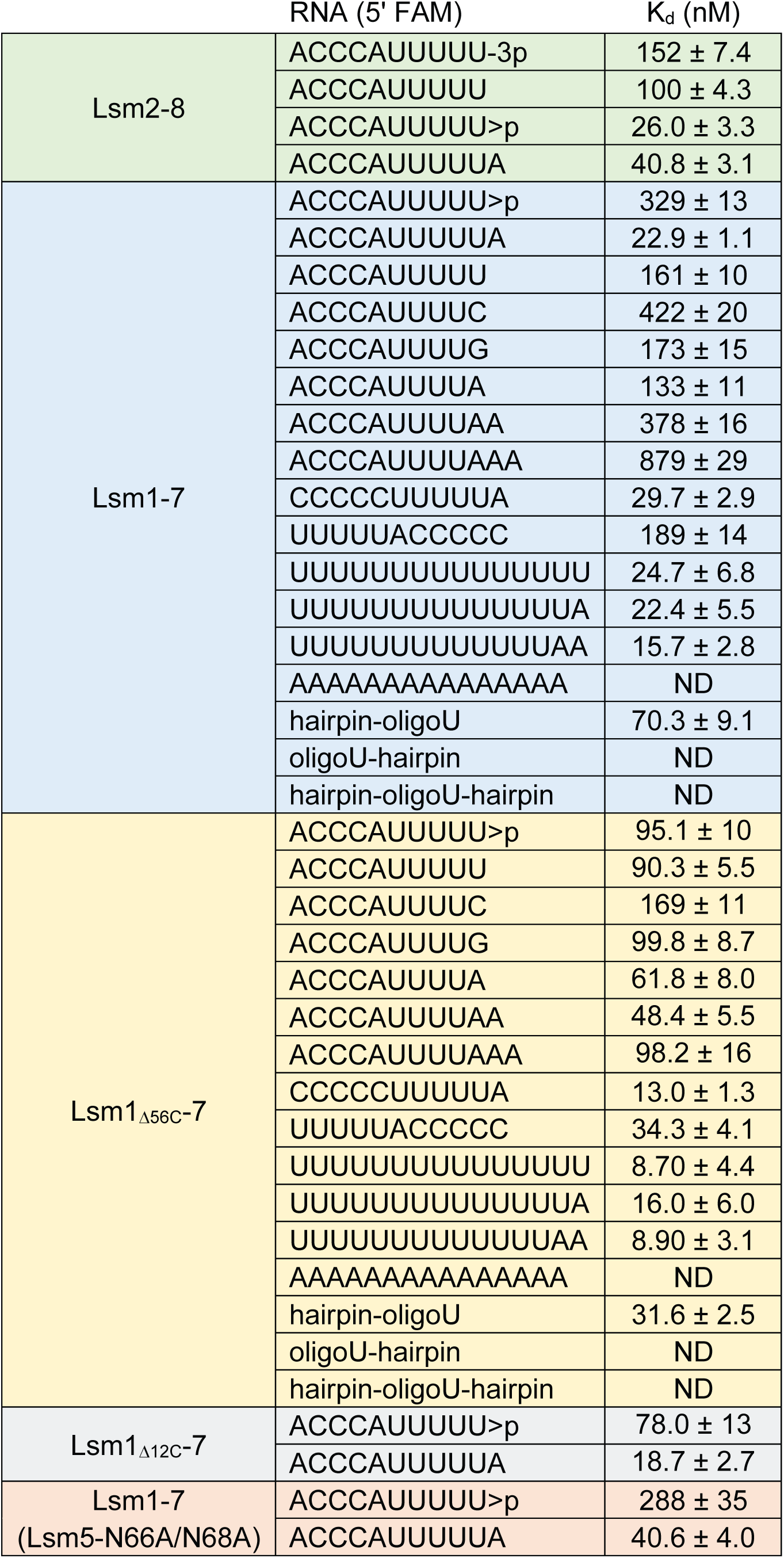
Summary of *in vitro* binding affinity between the indicated RNAs and *S. pombe* Lsm rings. All values were determined by fluorescence polarization binding assays with 5’ fluorescein-labeled RNA at 1 nM and a broad concentration range of Lsm proteins. Binding constants were determined by non-linear regression with a constrained Hill coefficient of 1. Raw polarization data and determined binding curves are available as Supplementary Data File 1.

**Supplementary Figure 1.**
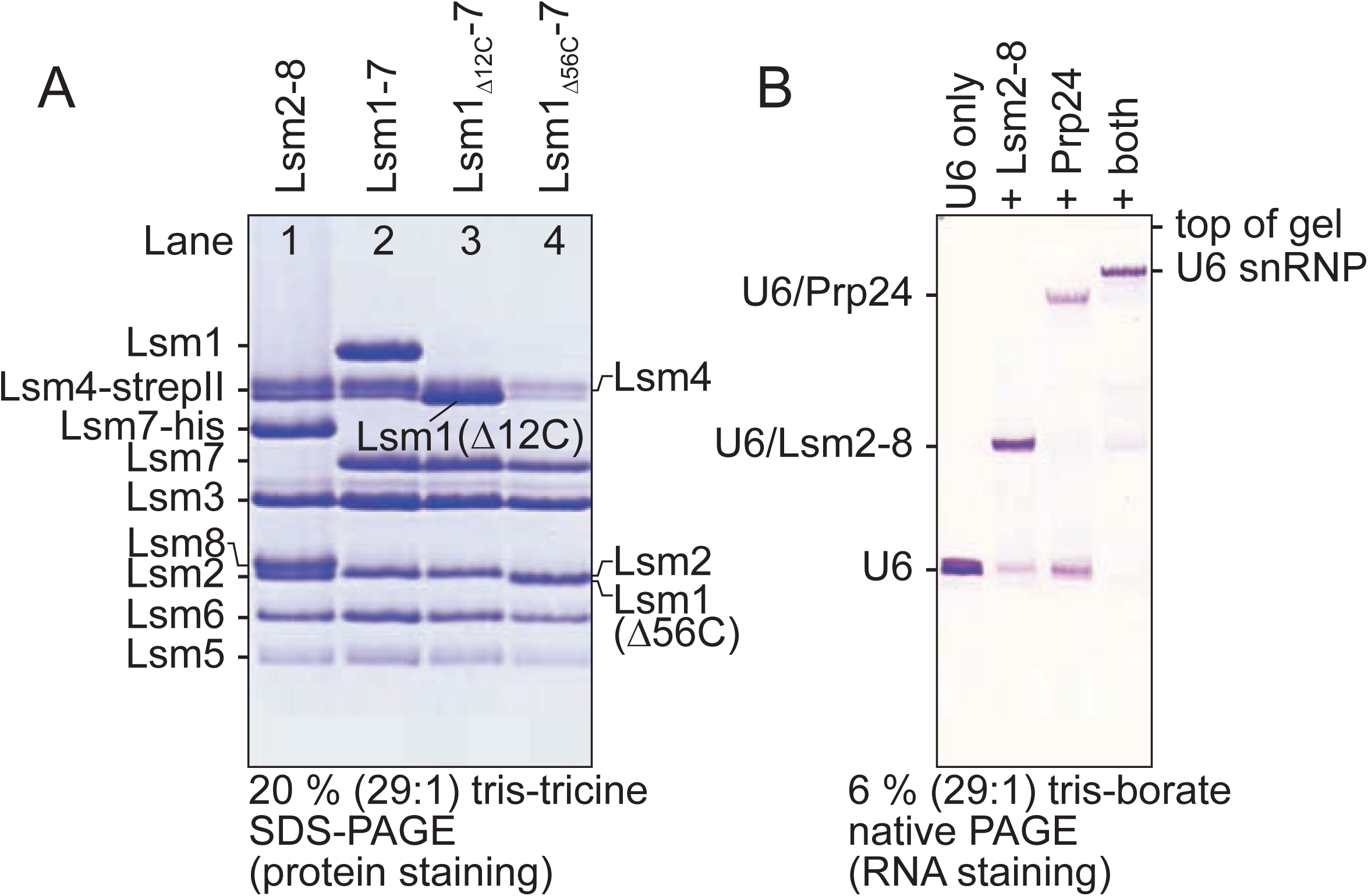
Production of recombinant *S. pombe* Lsm rings that can bind RNA and form macromolecular complexes. A) SDS-PAGE analysis of Lsm2-8 and Lsm1-7 proteins made by the pQLink system (36) in *E. coli* and purified with the use of multiple, cleavable affinity tags. Lsm4 appears as a very close doublet, either due to incomplete denaturation on SDS PAGE or partial proteolysis of its C-terminus which has no visible electron density in the crystal structures reported here and is predicted to be a region of low complexity. All masses for all Lsm proteins were confirmed by mass spectrometry and no proteolytic fragments of Lsm4 were detected by mass spectrometry. The Lsm7 subunit in Lsm2-8 (lane 1) retains a C-terminal oligohistidine tag that is not present in the final Lsm1-7 samples (lanes 2-4). B) *S. pombe* Lsm2-8 can form recombinant U6 snRNPs after addition of Prp24 and U6 snRNA.

**Supplementary Figure 2.**
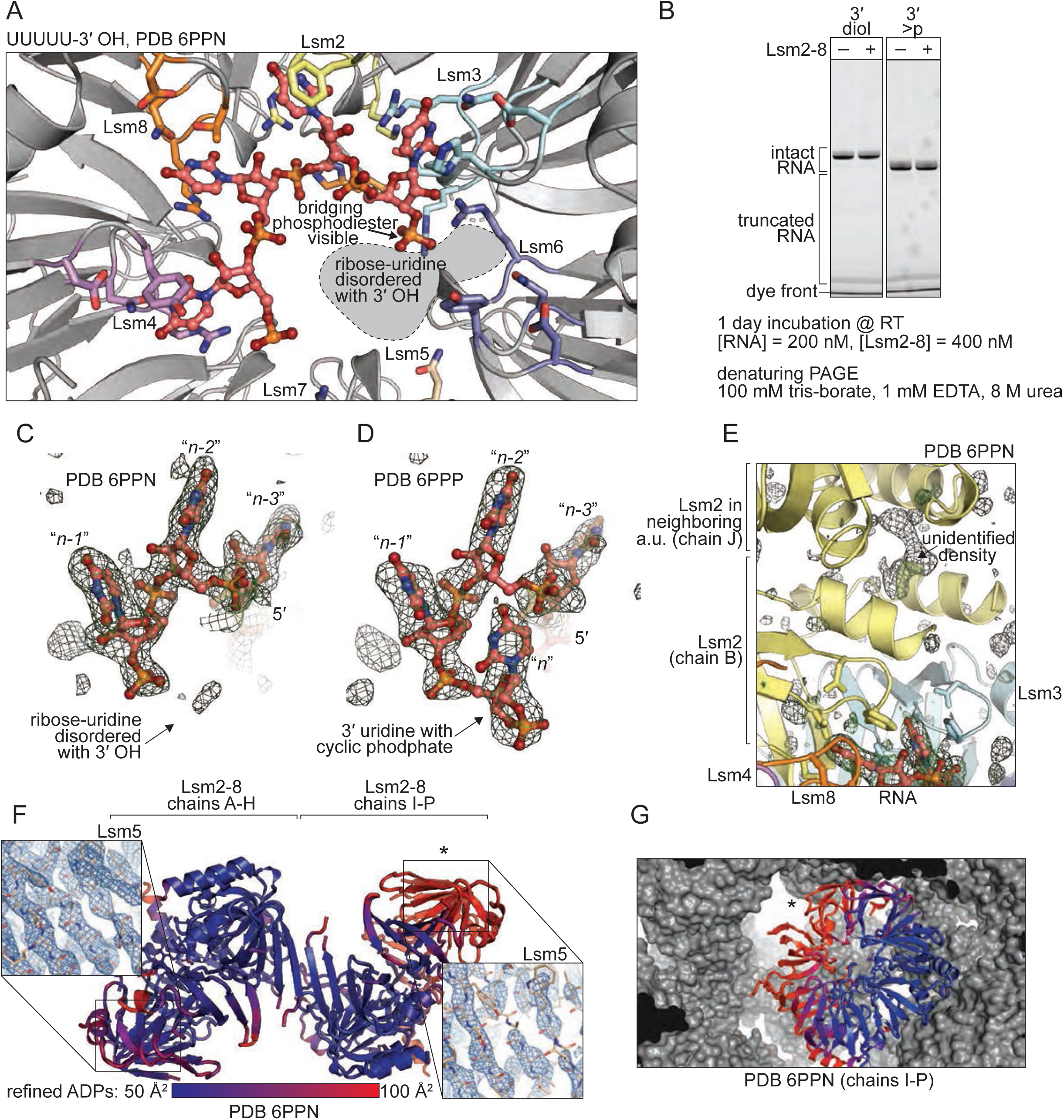
Structural details of *S. pombe* Lsm2-8 with RNA. A) Overview of unprocessed pentauridylate bound to Lsm2-8. Only the bridging phosphodiester between the last and penultimate uridine nucleotides is clearly visible in the electron density maps. B) Denaturing urea PAGE gels showing that prolonged incubation with Lsm2-8 does not result in cleavage and shortening of diol-terminated RNA, and as such the observed terminal phosphate in panel A is not likely to be a product of hydrolysis. C,D) Simulated annealing omit maps for RNA bound to Lsm2-8. Depicted maps are of form *m*F_o_-*D*F_c_, and are contoured at 1 r.m.s.d. E) Crystal packing of Lsm2-8 complexes in space group *P*2_1_2_1_2 is bridged by an unidentified buffer component that is putatively a contaminant within the pentaerythritol propoxylate (5/4 PO/OH) precipitant used for crystal growth. The depicted map is identical to that in panels C and D. F) Two Lsm2-8 complexes are present in the crystallographic asymmetric unit, which vary significantly in their local atomic displacement parameters (colored blue to red), especially around the Lsm5 subunits. Representative local 2*m*F_o_-*D*F_c_ maps at 1 r.m.s.d. are shown in the inserts. G) The poor density in one of the Lsm2-8 rings can be attributed to a lack of substantial crystal packing contacts in one copy of Lsm2-8. An asterisk denotes the same regions of poor main chain density in panels F and G.

**Supplementary Figure 3.**
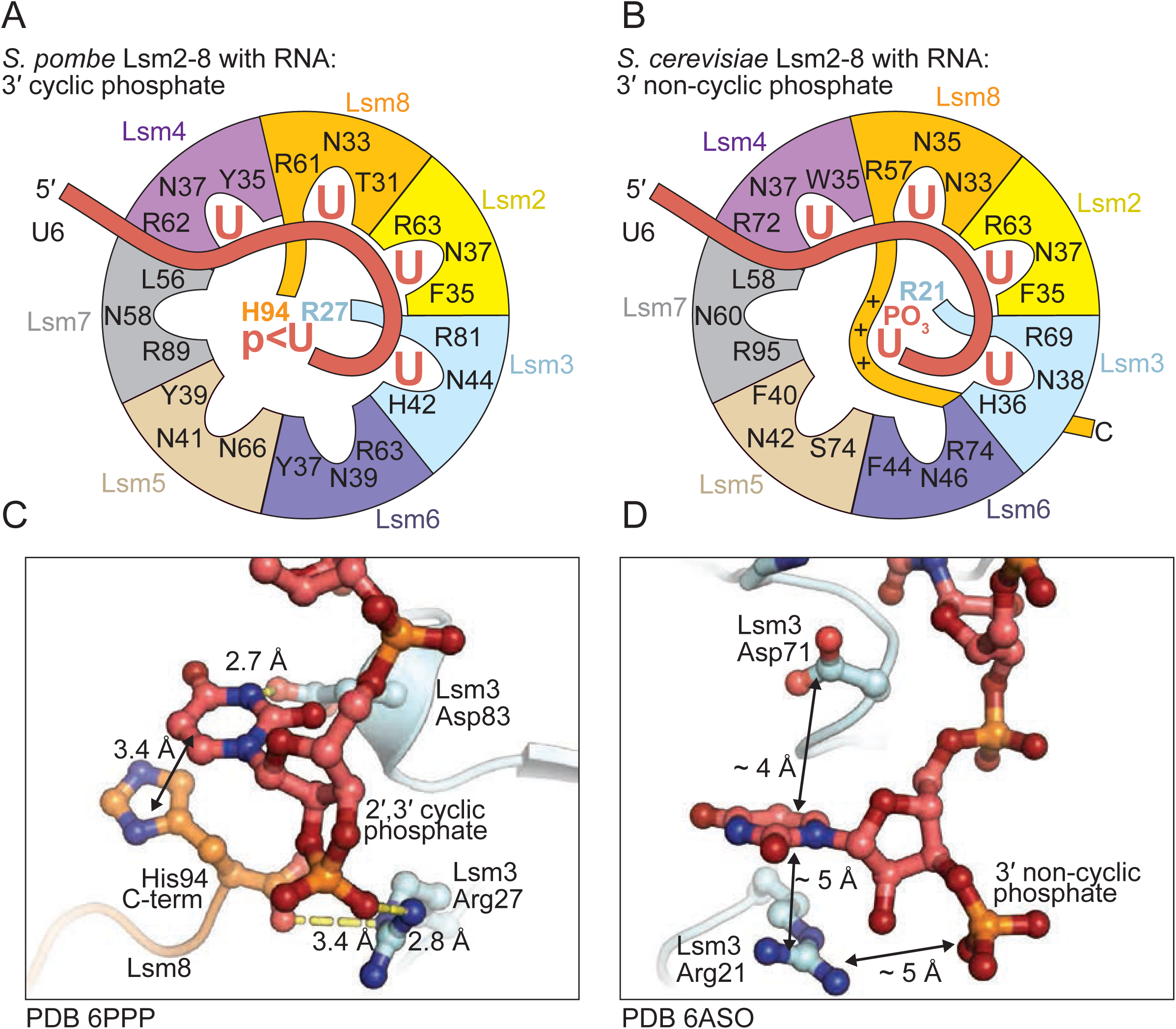
Comparison of how 3’ processed U6 RNA is recognized by *S. cerevisiae* and *S. pombe* Lsm2-8. A) *S. pombe* Lsm2-8 binds the 3′ uridine cyclic phosphate in the middle of the ring through non-Sm contacts with Lsm8 and Lsm3. B) The C-terminal tail of Lsm8 in *S. cerevisiae* is elongated and extends past the terminal uridine-phosphate. In lieu of direct interaction with 3′ uridine-phosphate, the tail of Lsm8 in *S. cerevisiae* interacts with the phosphate through long range electrostatics. C,D) Detailed comparisons of how the cyclic and non-cyclic phosphates are coordinated by the *S. pombe* and *S. cerevisiae* Lsm2-8 rings, respectively. Lsm3-Arg27 directly coordinates the cyclic phosphate in *S. pombe*.

**Supplementary Figure 4.**
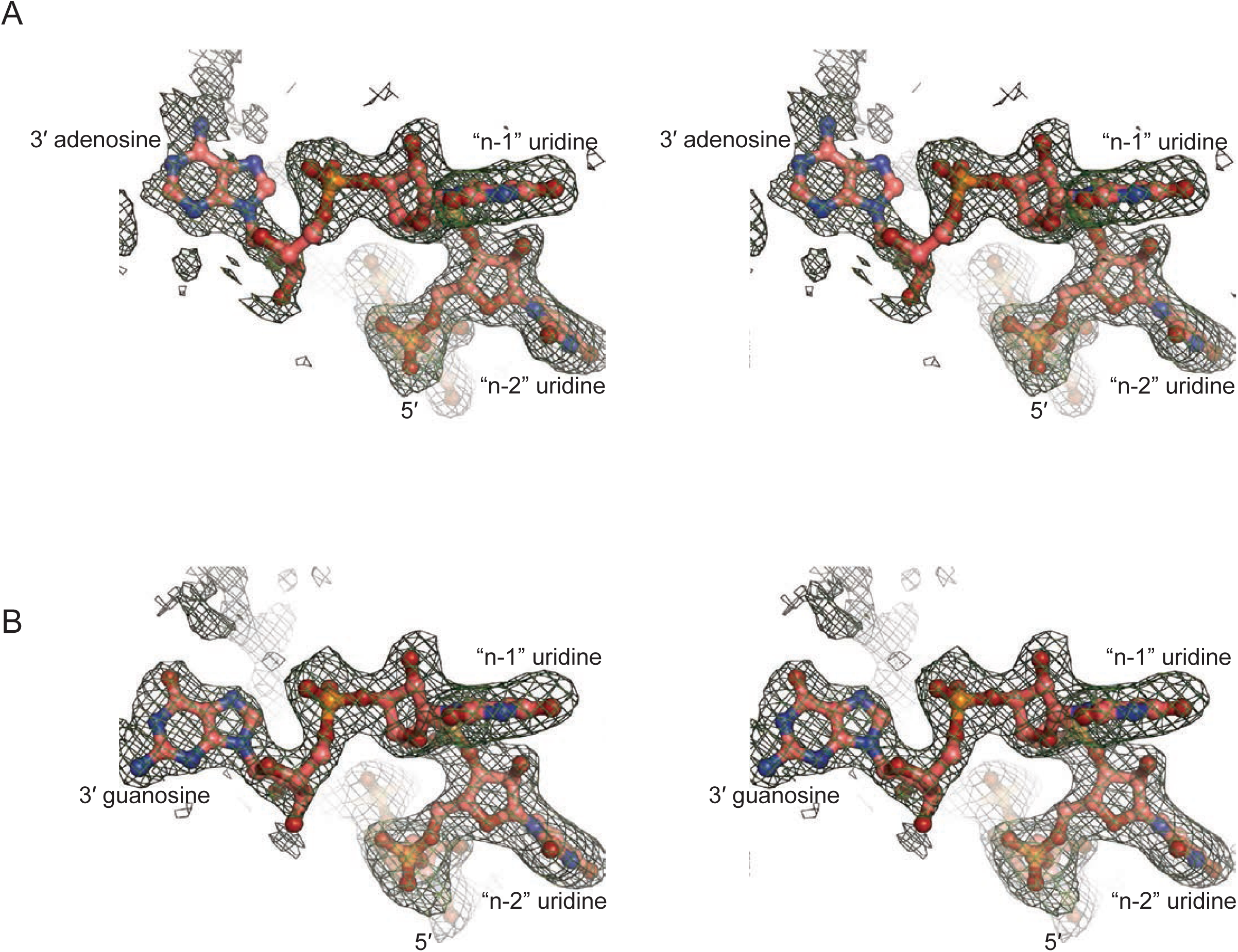
Simulated annealing omit maps depicting density for the 3′ purine residues bound by Lsm1-7. A) Cross-eyed stereo electron density map for adenosine terminated RNA. B) Map for guanosine terminated RNA. All maps are calculated and depicted similar to those in Supplementary Figure 2.

**Supplementary Figure 5.**
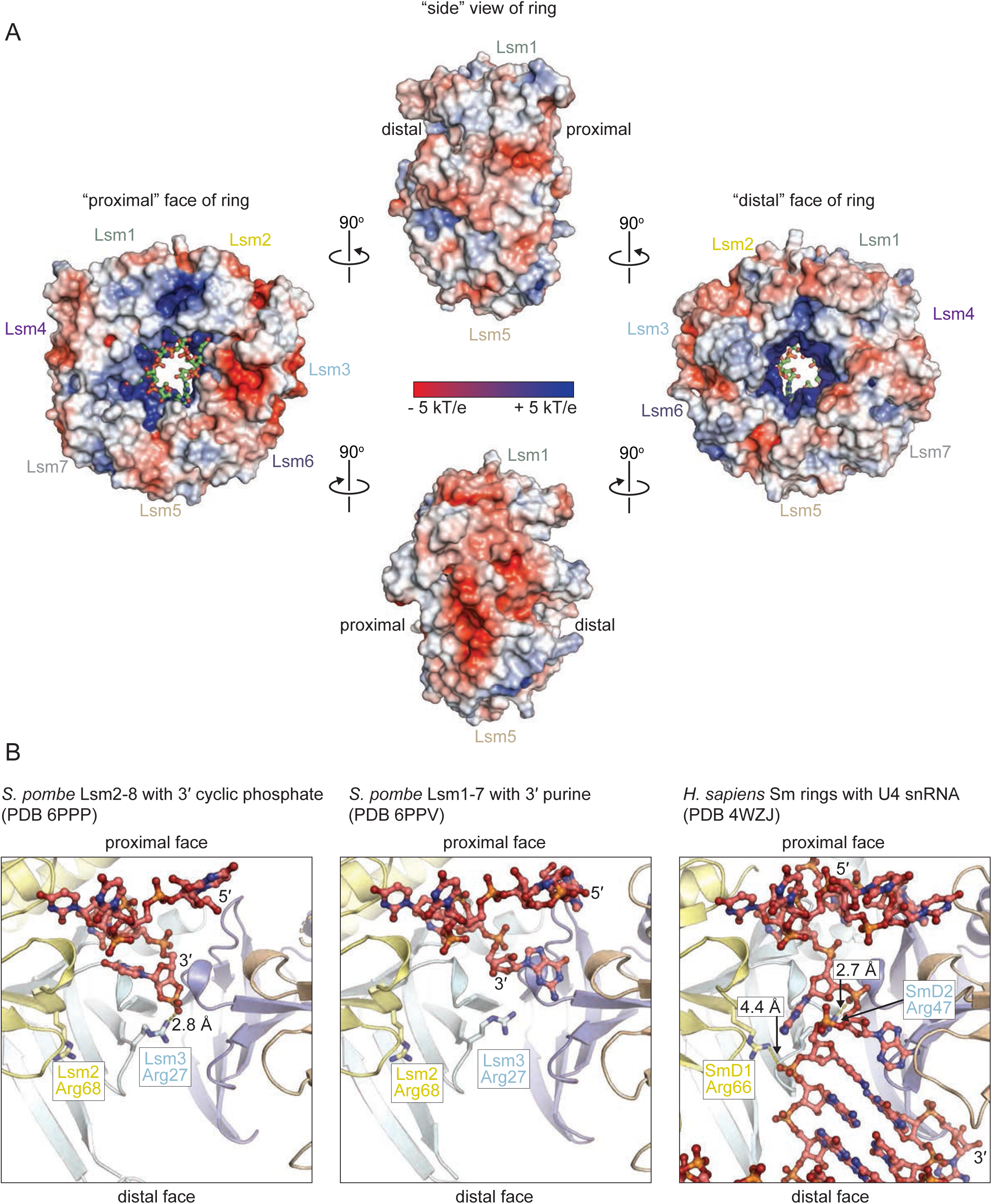
Surface properties of Lsm rings. A) Electrostatic surface analysis of *S. pombe* Lsm1_Δ56C_-7 bound to AUUUUG (PDB 6PPV). RNA is depicted in green for clarity. B) Comparison of the distal RNA entry sites of Lsm rings and the homologous Sm ring. Conserved residues in the Lsm and Sm rings are observed to interact with “threaded” RNA in the human U4 snRNP.

